# Lineage-Specific Epigenomic and Genomic Activation of the Oncogene HNF4A Promotes Gastrointestinal Adenocarcinomas

**DOI:** 10.1101/812149

**Authors:** Jian Pan, Tiago C. Silva, Nicole Gull, Qian Yang, Jasmine Plummer, Stephanie Chen, Kenji Daigo, Takao Hamakubo, Sigal Gery, Ling-Wen Ding, Yan-Yi Jiang, Shao-Yan Hu, Li-Yan Xu, En-Min Li, Yanbing Ding, Samuel J. Klempner, Benjamin P. Berman, H. Phillip Koeffler, De-Chen Lin

## Abstract

**Backgrounds:** Gastrointestinal adenocarcinomas (GIACs) of the tubular GI tract including esophagus, stomach, colon and rectum comprise most GI cancers and share a spectrum of genomic features. However, the unified epigenomic changes specific to GIACs are less well-characterized.We applied mathematical algorithms to large-scale DNA methylome and transcriptome profiles to reconstruct transcription factor (TF) networks using 907 GIAC samples from The Cancer Genome Atlas (TCGA). Complementary epigenomic technologies were performed to investigate HNF4A activation, including Circularized Chromosome Conformation Capture (4C), Chromatin immunoprecipitation (ChIP) sequencing, Whole Genome Bisulfite Sequencing (WGBS), and Assay for Transposase-Accessible Chromatin (ATAC) sequencing. In vitro and in vivo cellular phenotypical assays were conducted to study HNF4A functions.

**Results:** We identified a list of functionally hyperactive master regulator (MR)TFs shared across different GIACs. As the top candidate, HNF4A exhibited prominent genomic and epigenomic activation in a GIAC-specific manner. We further characterized a complex interplay between HNF4A promoter and three distal enhancer elements, which was coordinated by GIAC-specific MRTFs including ELF3, GATA4, GATA6 and KLF5. HNF4A also self-regulated its own promoter and enhancers. Functionally, HNF4A promoted cancer proliferation and survival by transcriptionally activating many downstream targets including HNF1A and factors of Interleukin signaling in a lineage-specific manner.

**Conclusion:** We use a large cohort of patient samples and an unbiased mathematical approach to highlight lineage-specific oncogenic MRTFs, which provide new insights into the GIAC-specific gene regulatory networks, and identify potential therapeutic strategies against these common cancers.

## Background

Tubular gastrointestinal adenocarcinomas (GIACs) including esophageal (EAC), stomach (STAD), and colorectal adenocarcinomas (COAD) arise from columnar epithelium and share similar endodermal cell-of-origin. GIACs represent one of the most common and aggressive tumor types, affecting over 3.5 million worldwide[1], and causing 1.4 million deaths[1] in the year of 2018 alone. At the genomic level, GIACs notably exhibit a spectrum of both common and unique characteristics, and can be subtyped accordingly into Epstein-Barr virus (EBV)-positive (STAD-specific), chromosomal instability (CIN), microsatellite instability (MSI) and genomically stable (GS)[2–4]. Some of these genomic features have important clinical implications. For example, EBV and MSI tumors showed increased sensitivity to immune checkpoint blockade[5, 6].

Large-scale genomic analyses[7–9] have established genetic landscapes of all three major GIAC types (EAC, STAD and COAD). Very recently, TCGA consortium conducted a comparative analysis and characterized shared genomic features across these GI cancers[4]. Compared with non-GIAC tumors, unique genomic changes in GIACs included amplification of lineage-specific transcription factors (e.g., *CDX2, KLF5* and *GATA4/6*) and mutations and deletions of *APC* and *SOX9*. Interestingly, this same study also noted that a significant fraction of pancreatic adenocarcinomas (PAAD) exhibited a high-degree molecular similarity with STAD (particularly the CIN and GS subtypes) based on consensus clustering results. Indeed, other individual reports also found that several key oncogenes (such as *GATA6* and *KLF5*[10–12]) are similarly essential for the proliferation and viability of both PAAD and other GIAC cancers.

Relative to genomic alterations, the epigenomic changes that are either shared or private in GIACs are less well-characterized. Most knowledge is derived from DNA methylation array data (Infinium HM450) which has identified the CpG island methylator phenotype (CIMP) uniquely in EBV+ and some MSI+ GIACs. Further integration with RNA profiling suggests that there conceivably exist GIAC-specific gene regulatory networks which contribute to the phenotypes and biological processes shared among GI cancers[7–9]. However, such gene expression networks and particularly their enhancer regions and upstream regulators have not been well elucidated in a Pan-GIAC manner.

Genome-wide DNA methylation profiling has shown that enhancer de-methylation represents the most dynamic change that occurs during neoplastic development[13–16]. Our earlier work demonstrated that this form of de-methylation could be used to identify cancer-specific enhancers which were strongly enriched for specific transcription factor binding sites (TFBSs)[14]. Recently, we have developed a novel computational algorithm, ELMER (Enhancer Linking by Methylation/Expression Relationships), to identify and exploit systematically these TFBS-associated methylation changes in cancer[17, 18]. Briefly, the approach of ELMER is to utilize TFBS methylation changes as key nodes within the larger gene regulatory network, and correlate with gene expression to infer both the upstream (master regulator TFs, MRTFs) and downstream (target genes) links for each TFBS. Specifically, when a group of enhancers is coordinately altered in a specific sample subset, this is often the result of an altered upstream MRTF in the gene regulatory network. To identify MRTFs involved in setting the tumor-specific DNA methylation patterns, ELMER correlates DNA methylation occurring within binding sites for specific MRTFs (inferred by specific TF binding motifs), with altered expression of the corresponding MRTF[17, 18].

Considering the molecular similarities among GIACs, we hypothesized that there exist GIAC-specific gene regulatory networks which are controlled by GIAC-specific MRTFs. Here, we addressed this hypothesis by applying ELMER to TCGA Pan-GIAC samples to explore whether different GIACs share functionally hyperactive MRTFs.

## Results

### Identification of hyperactive MRTFs in GIAC using ELMER

To identify master regulator TFs (MRTFs) with higher transcriptional activity in GIACs than adjacent normal samples, we analyzed paired transcriptome sequencing (RNA-Seq) and DNA methylation array data (Infinium HM450 Beadchip) from TCGA samples, including EAC (78 tumors *vs* 5 normal), STAD (338 tumors *vs* 2 normal), and COAD (389 tumors *vs* 21 normal). We additionally included PAAD (177 tumors *vs* 4 normal) since this cancer exhibits strong molecular similarity with a significant proportion of STAD, as introduced earlier[4]. The ELMER method was applied to compare tumor and normal specimens in each cancer type to identify MRTFs with higher activity in tumor samples (Fig. 1A). We further contrasted EAC *vs* ESCC tumors as they are biologically distinct subtypes from the same organ, serving as ideal analytical controls to discover EAC-specific networks. As a result, ELMER identified a total of 24 MRTFs (FDR q value <0.05), and notably, 10 of them were shared in at least two different comparisons (Fig. 1B). This result suggests that different GIACs indeed exhibit notable similarity in terms of their hyperactive MRTFs, which likely control similar downstream transcriptional programs. Some of these MRTFs are well-established GIAC specific oncogenes, such as *CDX2*[19, 20]*, GATA6*[10, 21]*, KLF5*[10, 11] *and HNF1A*[22], highlighting that ELMER is capable of identifying cancer-specific MRTFs.

**Figure 1.**
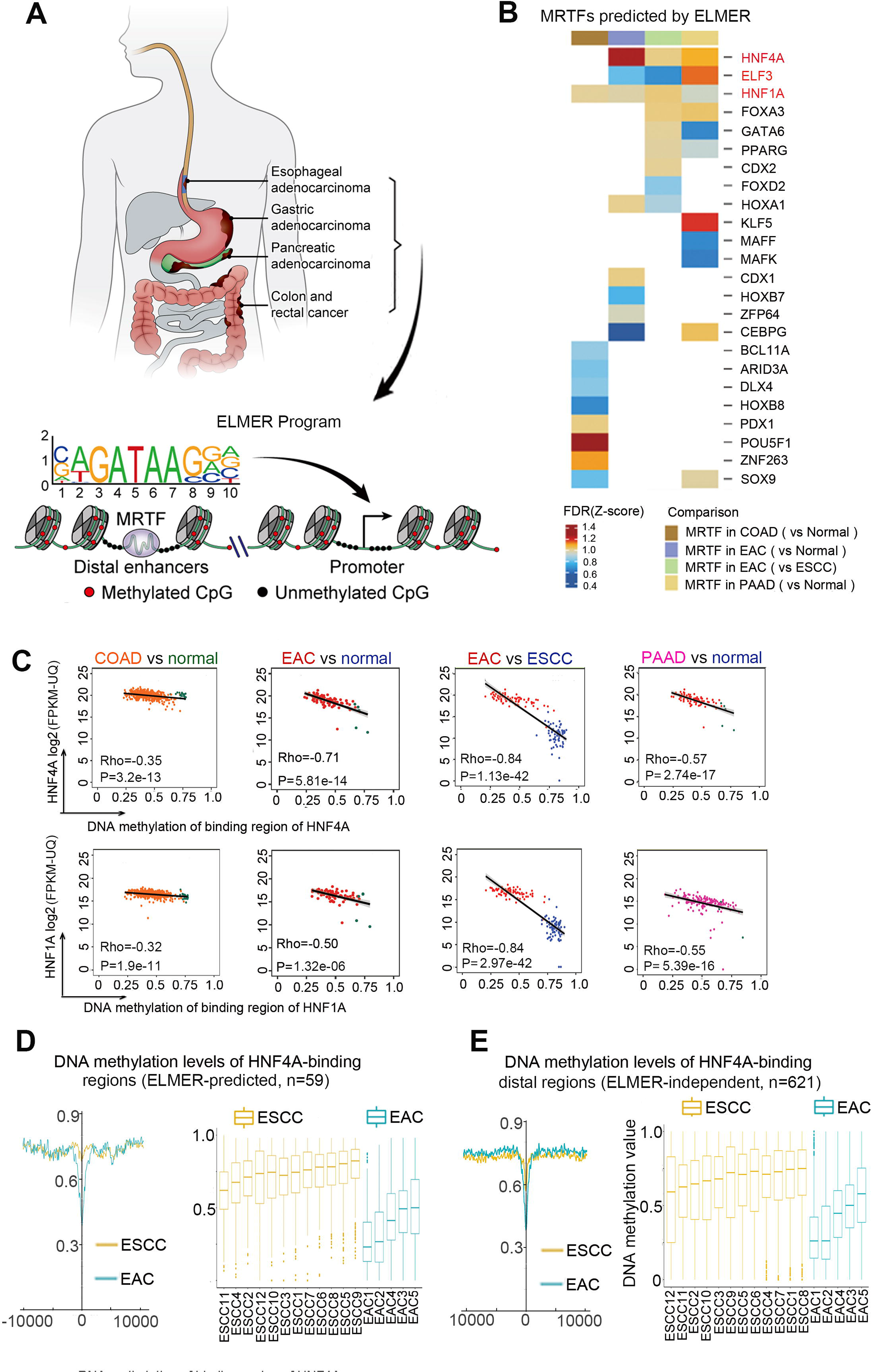
Identification of hyperactive MRTFs in GIAC using ELMER. (A) Schematic of the study design, showing tumor anatomical locations (top) and the ELMER approach for identifying Master Regulator Transcription Factors (MRTFs, bottom). (B) ELMER analysis was applied in each type of GIAC. A color is shown in each cell where the MRTF was identified (q < 0.05), and the color represents the p-value normalized for each GIAC (Z-score). (C) Scatter plots of top candidate MRTFs (HNF4A and HNF1A), showing the average DNA methylation level of predicted MRTF-binding regions (X axis) vs. the mRNA expression of the MRTF (Y axis) in each sample (dot). (D-E) Plots showing average WGBS methylation levels of predicted HNF4A binding sites, either averaged across all samples of the same subtype to show spatial pattern (left) or for each sample individually (right). (D) shows ELMER-predicted HNF4A binding sites, and (E) shows ESCA-specific ATAC-Seq peaks overlapping an HNF4A binding motif.

Since this work was aimed to discover hyperactive MRTFs shared by different GIACs, we focused on the only three candidates (HNF4A, ELF3 and HNF1A) which were identified in at least three comparisons (Figs. 1B). Among them, HNF4A was particularly interesting because: i) it had the lowest FDR q value among all 24 MRTFs; ii) it has been shown as an upstream regulator of HNF1A (another top-ranked MRTF) in hepatocytes[23], which we functionally validated in different types of GIAC cells (**Supplementary Fig. 1A**). Specifically, knockdown of HNF4A decreased the expression of HNF1A while the opposite was not observed (**Supplementary Figs. 1B-C**). Moreover, HNF4A directly occupied the promoter region of HNF1A in several different GIAC cell lines (**Supplementary Fig. 1A**). As shown in the scatter plots from each type of GIAC (Fig. 1C), HNF4A-high tumors had significantly lower DNA methylation in predicted HNF4A binding regions than either HNF4A-low tumors or normal tissues. The ELMER analysis was based on the Infinium HM450 design, which covers less than 5% of all enhancers[24]. To validate the ELMER results, we generated whole genome bisulfite sequencing (WGBS) data from an independent cohort of 5 EAC and 12 ESCC samples, which covered an average of 97.29% of all CpGs in the genome with at least 5 sequencing reads. We first determined the DNA methylation levels of ELMER-predicted HNF4A-binding regions and confirmed that indeed they were lower in EAC than ESCC WGBS samples (Fig. 1D). To further validate the de-methylation of HNF4A binding regions genome-wide, we next performed a completely independent analysis without using the ELMER results. Briefly, we analyzed methylation within ESCA-specific open chromatin regions from TCGA ATAC-Seq (12 ESCC and 6 EAC samples)[25] which contained HNF4A binding motif sequences (see Methods). As expected, these predicted HNF4A binding sites were consistently hypomethylated in EAC samples than ESCC samples (Fig. 1E). Moreover, the DNA methylation difference was much more pronounced in distal regions than promoters (**Supplementary Fig. 1E**), consistent with the distal location of binding sites from HNF4A ChIP-seq in GIAC cells (**Supplementary Fig. 1D**).

### Unique genomic and epigenomic activation of HNF4A in GIACs

The above results suggest that HNF4A is highly activated in GIACs. Indeed, Pan-Cancer RNA-Seq data from TCGA showed that the expression of HNF4A was over-expression in only 7 out of 32 cancer types and top 4 were all GIACs (Fig. 2A). Although HNF4A was expressed comparatively high in hepatocellular, cholangio and kidney cancers, it was down-regulated comparing with the correspondent nonmalignant tissues. In the remaining 19 tumor types, HNF4A was undetectable in either nonmalignant or cancerous samples (Fig. 2A). HNF4A protein was then interrogated using the immunohistochemistry (IHC) results from The Human Protein Atlas[26, 27], and consistently, its protein level was most abundant in GIACs across all cancer types, displaying a prominent lineage-specific expression pattern (Fig. 2B). We next performed IHC staining on three cohorts of internal GIAC samples, and confirmed that HNF4A protein was significantly up-regulated in EAC, PAAD and COAD tumors compared with adjacent normal samples (Fig. 2C). Moreover, higher HNF4A expression was significantly associated with worse overall survival in both EAC and COAD patients (Fig. 2D). Searching for the mechanisms underlying the GIAC-specific over-expression of HNF4A, we found that with the exception of uterine tumors, the *HNF4A* gene locus was specifically amplified in 7-9% of different GIAC types, and had low-level copy number gain in a majority of GIAC samples (Figs. 3A-B). Expectedly, *HNF4A*-amplified and copy number gain tumors had higher HNF4A mRNA level than non-amplified tumors (Fig. 3C). However, we reasoned that copy number gain of *HNF4A* did not completely account for its mRNA upregulation, and epigenetic changes were also involved. This was because gene amplification only increased its mRNA log2 FPKM level by 2-4 (Fig. 3C), whereas its log2 FPKM was elevated by at least 10 compared with most non-GIAC samples (Fig. 2A). To explore the possibility of epigenetic upregulation, we queried TCGA ATAC-Seq data (Fig. 3D) and observed that indeed the *HNF4A* promoter was markedly more accessible in GIAC samples than other cancer types (such as squamous tumors). Furthermore, several distal *HNF4A* intronic regions also exhibited higher accessibility (Fig. 3D). We aligned these ATAC-Seq tracks with in-house WGBS data and found that GIAC-specific open chromatin regions were concordantly de-methylated in EACs but fully methylated in ESCCs (Red arrows, Fig. 3D), suggesting that these are cis-regulatory elements potentially contributing to the transcriptional activation of HNF4A in GIAC in a lineage-specific manner.

**Figure 2.**
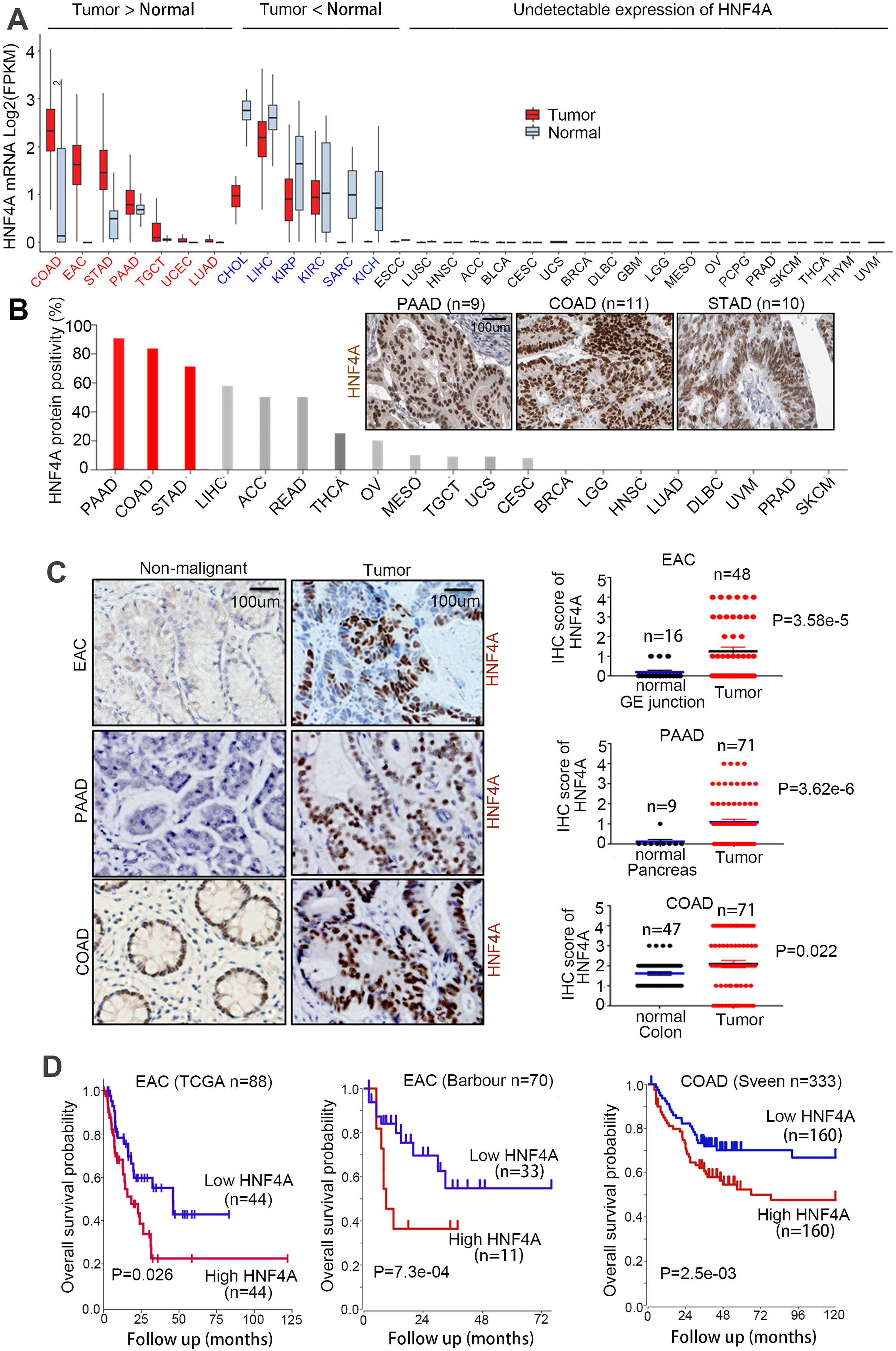
HNF4A is specifically and highly expressed in GIACs. (A) Expression of HNF4A in RNA-Seq data from TCGA Pan-Cancer samples. Each box represents a cancer type (TCGA abbreviations can be found at Supplemental Table 3. (B) HNF4A protein expression was interrogated using the IHC data from Human Protein Atlas project, and GIACs are highlighted in red. Representative IHC photos are shown as inserted panels. (C) IHC staining on 3 GIAC cohorts of internal samples. (D) Overall survival analyses of HNF4A expression in both EAC and COAD patients, from TCGA and other named studies.

**Figure 3.**
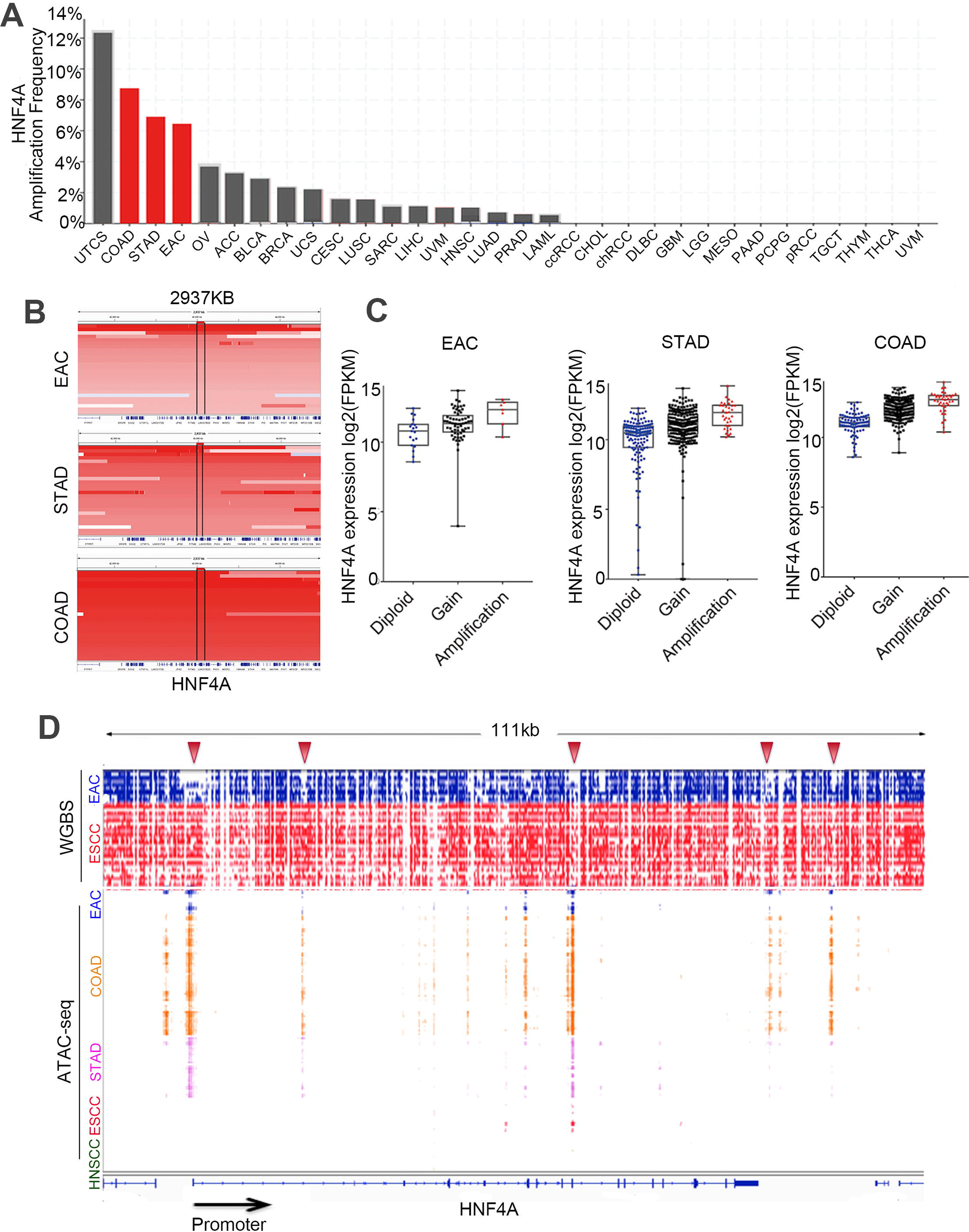
Genomic and epigenomic activation of HNF4A is specific to GIACs. (A) *HNF4A* gene amplification in SNP-array data from TCGA Pan-Cancer samples, with GIACs highlighted in red. (B) IGV (Integrative Genomics Viewer) plots of *HNF4A*-amplified GIACs samples. (C) mRNA level of HNF4A in GIAC samples with different *HNF4A* genomic status. Copy number *gain* refers to samples with between 2 and 4 copies of the *HNF4A* locus, and *amplification* refers to those with greater than 4 copies. (D) IGV plots showing *HNF4A* locus using both in-house WGBS samples (EAC=5, ESCC=12) and TCGA ATAC-Seq samples (EAC=6, COAD=38, STAD=21, ESCC=12, HNSCC=9). Arrows show several regions that are specifically accessible and demethylated in GIAC tumor types.

### HNF4A specifically promotes the proliferation and survival of GIAC cells

The above results demonstrate that HNF4A is specifically over-expressed in Pan-GIAC samples because of both genomic and epigenomic alterations. We thus hypothesized that this is a *bona fide* MRTF playing a functional role in GIAC biology. Indeed, HNF4A was reported to be essential for the proliferation and metabolism of STAD cells[10, 28, 29]. Nevertheless, it is unclear whether HNF4A activity is functionally required for other GIAC types including EAC, PAAD and COAD. Moreover, its biological functions and associated molecular mechanisms in GIAC remain incompletely understood. To address these questions, we initially silenced HNF4A expression in cell lines from EAC, COAD and PAAD with high endogenous level of this MRTF. Cell proliferation and colony formation capacity were strongly and consistently suppressed in HNF4A-knockdown group compared with control cells (Figs. 4A-C). We also noted that HNF4A-depletion caused massive apoptosis in all types of GIAC cells (Fig. 4D**, Supplementary Figs. 2A,B**). We next asked whether excessive levels of HNF4A could promote GIAC proliferation in GIAC cells with low endogenous HNF4A. Indeed, its ectopic over-expression enhanced the proliferation and colony formation in EAC, COAD and PAAD cells (Figs. 4E-G).

**Figure 4.**
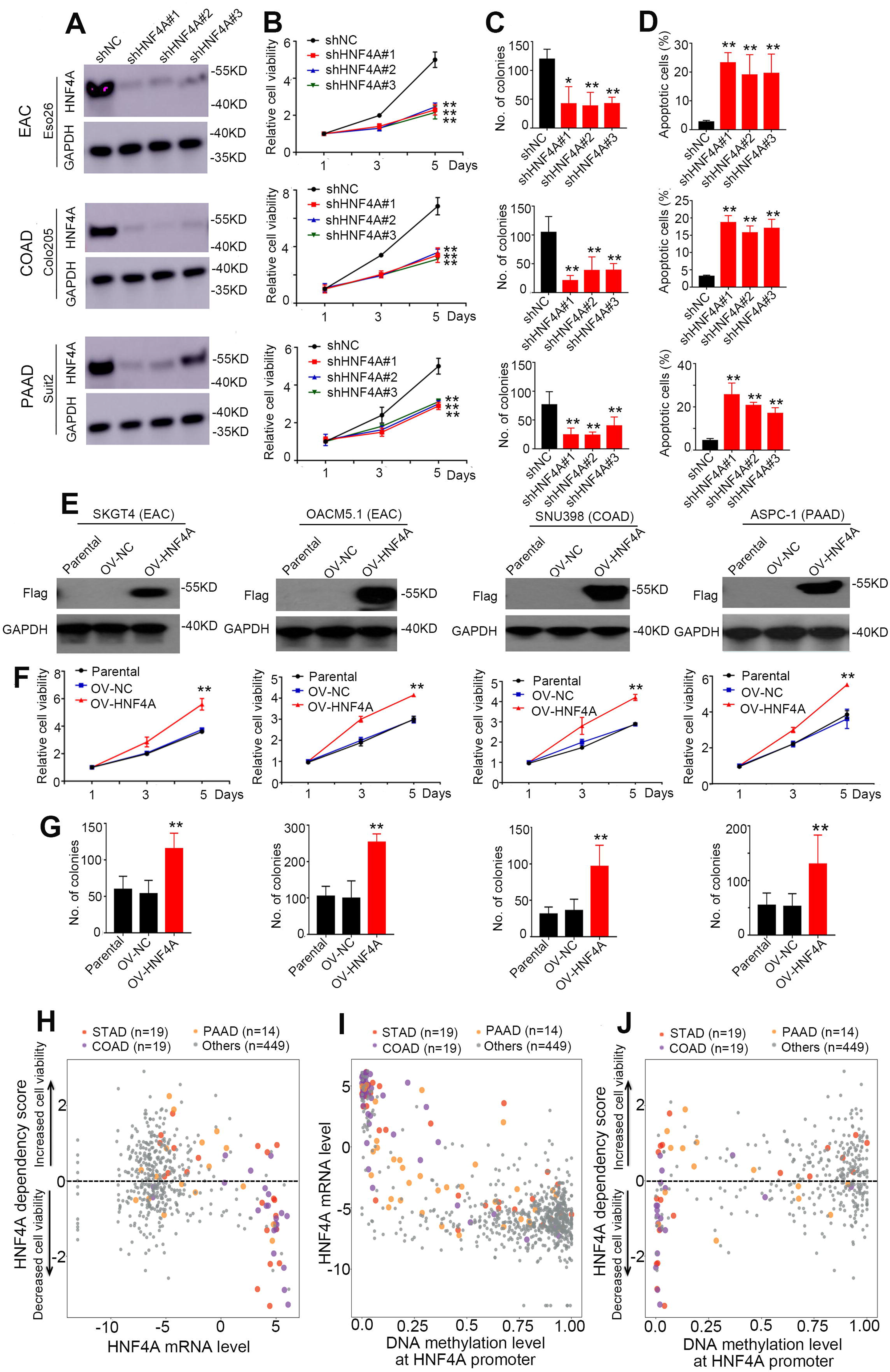
HNF4A specifically promotes the proliferation and survival of GIAC cells. In EAC (Eso26), COAD (Colo205) and PAAD (Suit2) cell lines, HNF4A expression was silenced by three different shRNAs and followed by Western Blotting assay (A), cell proliferation assay (B), colony formation assay (C), and apoptosis assays (D). In (E), HNF4A was ectopically expressed and validated by Western Blotting in EAC (SKGT4 and OACM5.1), COAD (SNU398) and PAAD (ASPC-1) cells. HNF4A-overexpressing cells were subjected to cell proliferation (F) and colony formation (G) assays (OV-NC, overexpression control; OV-HNF4A, overexpression of HNF4A). Data is presented as Mean ± s.d. *, P<0.05; **, P<0.01. (H) High-throughput shRNA screen in a pan-cancer analysis of 500 cell lines from CCLE and DepMap. Scatter plot showing the HNF4A shRNA dependency score vs. the expression of HNF4A. (I) Scatter plot showing the mRNA expression of HNF4A and the methylation of its promoter. (J) Scatter plot showing the HNF4A dependency score vs. the methylation of HNF4A promoter. Each dot represents one cell line, and GIAC cells are highlighted.

Considering that HNF4A is specifically activated in GIAC cancers, we next sought to test whether the functional significance of this MRTF exhibits similar GIAC specificity. Expectedly, HNF4A mRNA level was the highest in GIAC cells in a pan-cancer analysis of over 1,000 cell lines from CCLE (Cancer Cell Line Encyclopedia)[30](Fig. 4H). Strikingly, HNF4A was uniquely required for the viability of GIAC cells in the unbiased high-throughput shRNA screen (Fig. 4H). Consistent with our WGBS data, the HNF4A promoter was hypomethylated and correlated with high expression uniquely in GIAC cell lines in the CCLE dataset (Fig. 4I). We next tested whether its promoter methylation lever was similarly associated with its essentiality for cell viability. Indeed, cells with lower methylation of HNF4A promoter were more dependent on HNF4A for proliferation in a GIAC-specific manner (Fig. 4J). These results together identified HNF4A as a prominent lineage-specific oncogene in GIAC tumors.

### Targeting HNF4A suppresses growth of GIAC cancer cells and xenografts

Using a Dox-inducible shRNA system, we tested HNF4A function *in vivo* using an EAC-derived xenograft model (Fig. 5A). Both the growth and weight of the tumor xenografts were significantly inhibited by HNF4A-silencing (Figs. 5B-C). IHC analysis showed that Ki-67 expression (a marker of cell proliferation) was down-regulated in HNF4A-knockdown tumors (Fig. 5D). We next investigated the function of HNF4A by a small-molecule inhibitor (BI-6015[31]) developed against this MRTF. This antagonist binds to HNF4A with high affinity by forming hydrogen bonds with HNF4A Arg226 and Gly237 which occupies a hydrophobic pocket. This occupancy by BI-6015 inhibits the transcriptional function of HNF4A by preventing interaction with its ligand binding pocket. To test the effects of BI-6015 *in vitro*, we first conducted luciferase reporter assays using promoters from two direct HNF4A targets, *HNF1A* and *HNF4A* itself (direct binding at the HNF1A promoter is shown in **Supplementary Fig. 1A**, and autoregulation of the *HNF4A* promoter is shown below). BI-6015 strongly inhibited the reporter activities of both promoters, as well as the mRNA levels of both genes (Fig. 5E). In an *in vitro* proliferation assay, different GIAC cells from either EAC, PAAD or COAD were significantly more sensitive to this compound than non-GIAC cells (Figs. 5F-G), and that cell sensitivity to BI-6015 (measured by IC50) was negatively correlated with the expression of HNF4A (Fig. 5H), again confirming the on-target effect. The mouse xenograft assay showed that BI-6015 potently inhibited the growth of GIAC tumors but did not cause systematic toxicity (Fig. 5I**, Supplementary Figs. 3A,B**). Together, these results obtained from genetic and chemical approaches demonstrate that HNF4A functionally promotes the proliferation and survival specifically in GIAC cancer cells.

**Figure 5.**
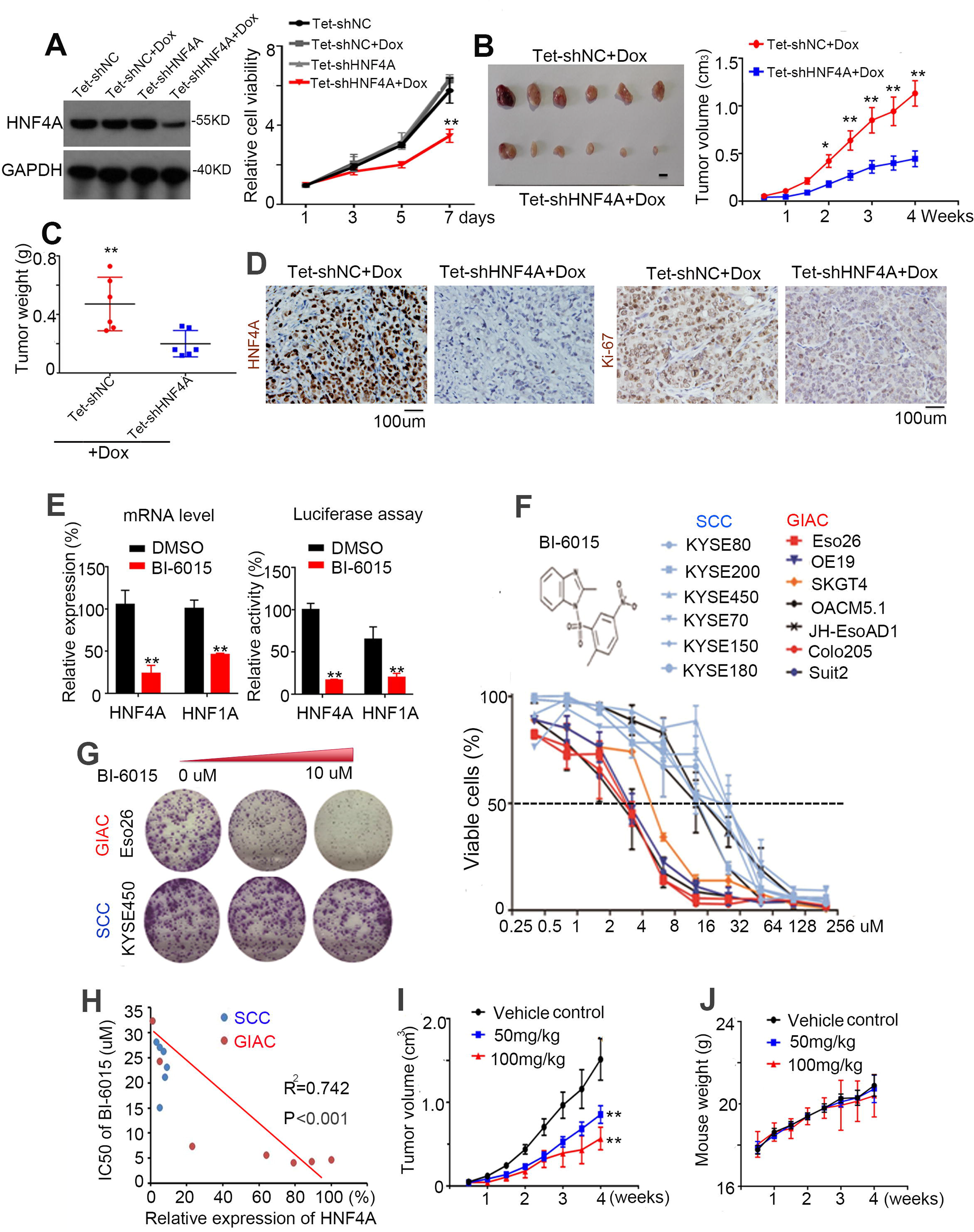
Targeting HNF4A suppresses growth of GIAC cancer cells and xenografts. (A) Western blotting assay (Left) validating the knockdown of HNF4A using Dox-inducible shRNA in Eso26 cells, which led to decreased cell proliferation in vitro (Right). (B) Tumor growth and (C) weight of the xenografts were significantly inhibited by HNF4A-silencing. (D) IHC staining of HNF4A and Ki-67 in xenograft samples. (E) qRT-PCR analysis (Left) and luciferase reporter assay (Right) validating the on-target effect of BI-6015. (F) Dose-response curves of GIAC and squamous cancer cell (SCC) lines treated with BI-6015 (72 hr). Data represent mean ± SD of three replicates. (G) Colony formation of Eso26 (EAC cell) and KYSE450 (SCC cell) treated with increasing dose of BI-6015. (H) Scatter plot showing the negative correlation between cell sensitivity (measured by IC50 of BI-6015) and HNF4A expression. Each dot represents one cell line. (I) Xenograft assay showed that BI-6015 inhibited the growth of GIAC tumors but did not affect the weight of the mice (J). Mean ± s.d. are shown. *, P<0.05; **, P<0.01.

### Upstream regulation of HNF4A transcription by GIAC MRTFs

To probe the mechanism underlying GIAC-unique epigenomic activation of HNF4A, we first performed circularized chromosome conformation capture (4C) sequencing to explore the interaction landscape of the *HNF4A* locus in EAC cells, using its promoter as the 4C-Seq bait (Viewpoint). We identified that all of the significant interactions (q < 0.001) were restricted to the 500kb window flanking the *HNF4A* promoter (Fig. 6A). Importantly, by cross-referencing H3K27Ac ChIP-Seq data generated in the matched sample, we found that the majority of interacting regions had positive H3K27Ac signals (Fig. 6B), suggesting their potential regulatory function. Based on the q value, we identified top 10 most significant interactions which overlapped with three H3K27Ac+ regions (referred to as E1-E3, Fig. 6C). To search for potential TFs occupying these regions, we performed motif enrichment analysis and found that top enriched motifs included the GATA family, ELF3 and HNF4A itself (Fig. 6D). This result was encouraging since GATA4/6 are known GIAC-specific MRTFs and ELF3 was the 2^nd^ most highly ranked MRTF identified earlier (Fig. 1B). Furthermore, the enrichment of HNF4A motif is also in keeping with the notion that self-regulation is an important and common property of many MRTFs in different cell types [32, 33]. To validate this motif analysis, we performed ChIP-Seq, wherein we additionally included KLF5 as our recent work identified it as an integral component of core regulatory circuitry (CRC) in EAC (Manuscript In Revision). Importantly, the ChIP-Seq results confirmed that the four regulatory regions (promoter region and E1-E3) were indeed occupied by GATA4, GATA6, ELF3, HNF4A and KLF5 in different types of GIAC cell lines (Fig. 6C**, Supplementary Fig. 4**). Notably, both E3 and promoter regions were co-occupied by all 5 MRTFs (Fig. 6C). To test the transcriptional activity of these regulatory elements, they were cloned individually into the luciferase reporter vector. All three enhancer elements showed robust reporter activities uniquely in GIAC cells but not in SCC cells (Fig. 6E). Moreover, silencing each of the 5 MRTFs inhibited the activities of promoter and E3 region. In addition, ELF3, KLF5 and HNF4A contributed to the activity of E1 and E2 (Fig. 6F). These results demonstrated strong and complex regulation of HNF4A transcription, which is GIAC-specific and cooperatively controlled by GATA4, GATA6, ELF3, KLF5 and HNF4A itself.

**Figure 6.**
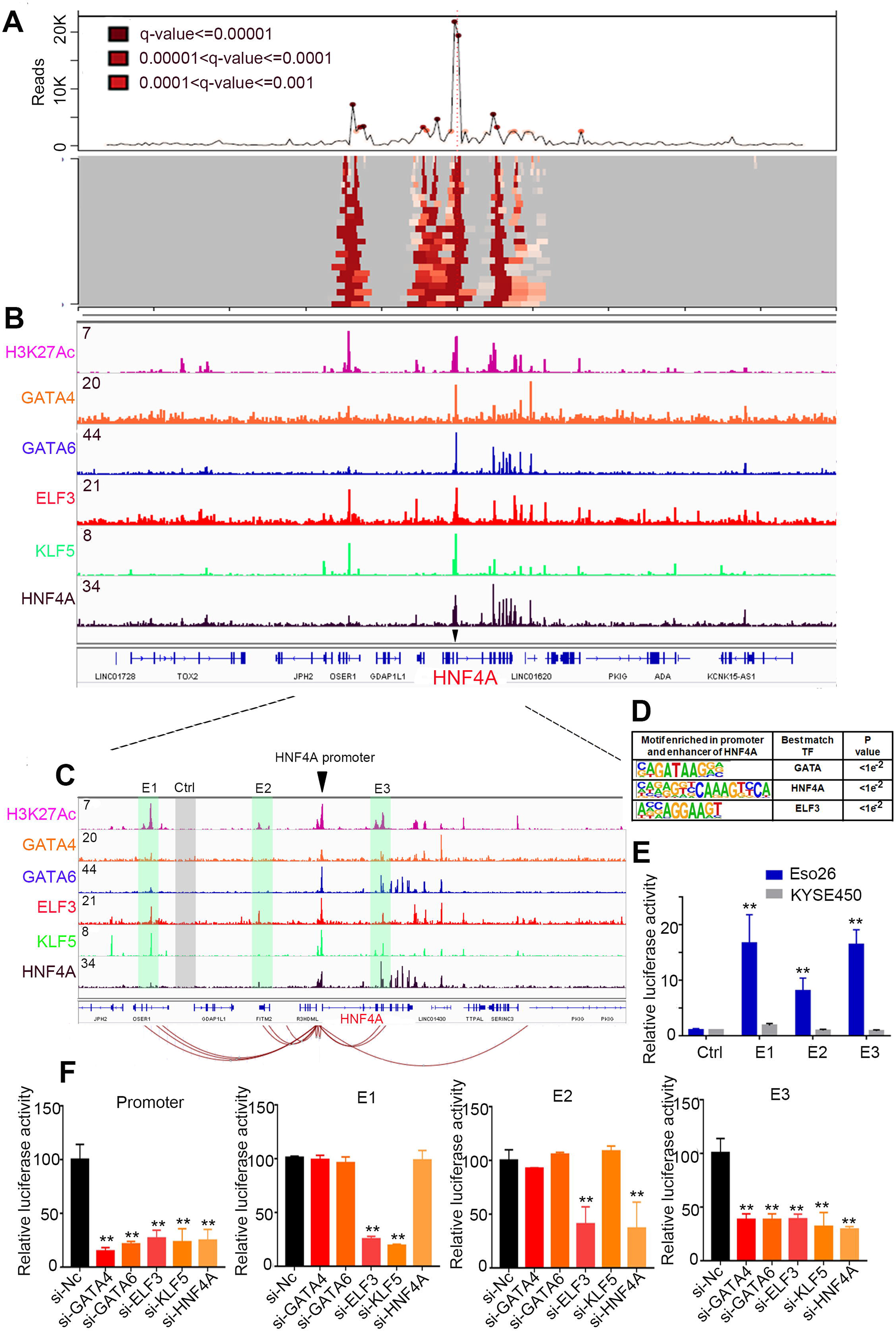
GIAC MRTFs occupy and regulate HNF4A promoter and enhancers. (A) 4C-Seq assay showing the long-range interactions anchored on HNF4A promoter in Eso26 cells. Deeper red color indicates higher interaction frequency. (B) ChIP-Seq profiles for H3K27Ac and indicated MRTFs at HNF4A loci in Eso26 cells. (C) Zoom in view of ChIP-Seq signals. Connecting lines showing the most significant interactions detected by 4C. Shaded regions showing three enhancers (E1-E3) and one negative control (Ctrl) region which were separately cloned into the luciferase reporter vector. (D) Motif enrichment analysis of promoter and E1-E3 regions. (E) Enhancer activity measured by luciferase assays in Eso26 and KYSE450 cells. (F) Enhancer and promoter activity measured by luciferase assays in Eso26 cells upon silencing each of the 5 MRTFs. Mean ± s.d. are shown. *, P<0.05; **, P<0.01.

Importantly, individually silencing each MRTF significantly down-regulated the expression of HNF4A at both mRNA and protein levels, confirming the regulation of HNF4A by these factors (Fig. 7A-B). Given the co-localization of these 5 MRTFs in HNF4A promoter and enhancers, we tested if there existed direct protein-protein interactions among these upstream factors. Immunoprecipitation (IP) analysis showed that GATA4, GATA6 and KLF5 formed protein complexes in GIAC cells (Fig. 7C**, Supplementary Fig. 5B**). While this protein complex did not contain HNF4A or ELF3, these two MRTFs might participate in the transcription co-operation through indirect mechanisms (indirect cooperativity, such as the “Billboard model” [34]). Considering that some of these MRTFs (e.g., GATA4/6 and KLF5) have been reported to have copy number gains in GIAC samples[10, 11], and our present work identified that *HNF4A* was also amplified specifically in GIACs, we next analyzed the genomic status of these MRTFs in TCGA dataset. The copy number gains of *GATA4/6* and *KLF5* were confirmed in both EAC and STAD cohorts (**Supplementary Fig. 5A**). Notably, the amplification of these MRTFs exhibited a mutually exclusive pattern (**Supplementary Fig. 5A**), strongly supporting our results that these MRTFs converge on HNF4A signaling, and thus redundant gain-of-function events are not required in the same GIAC samples.

**Figure 7.**
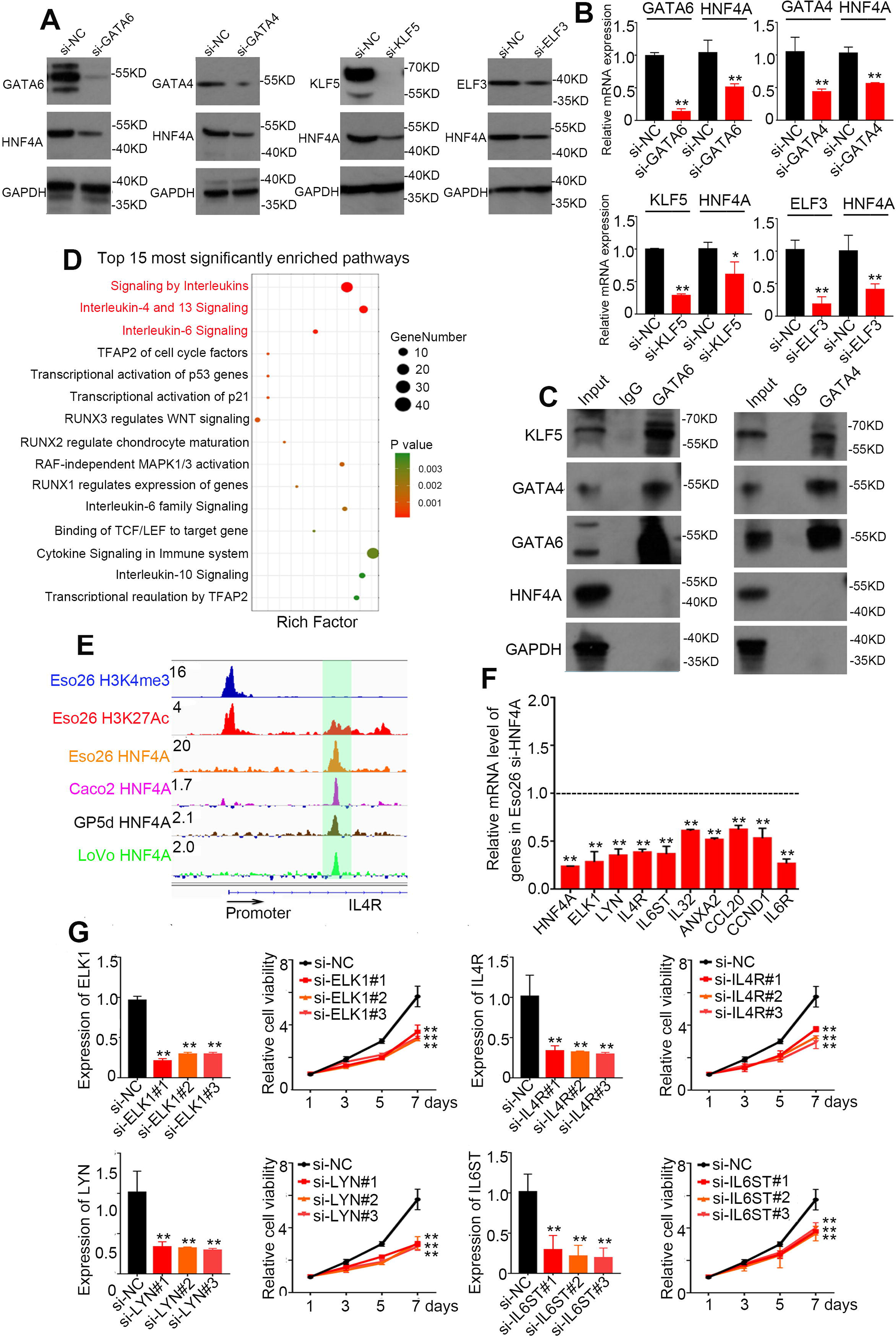
Upstream and downstream regulation of HNF4A in GIAC cells. (A) Western Blotting and (B) qRT-PCR assay upon each individual knockdown of indicated MRTFs in Eso26 cells. (C) Co-IP assay of GATA4, GATA6, and HNF4A in Eso26 cells. (D) Pathway enrichment analysis of the differentially expressed genes upon silencing of HNF4A in Eso26 cells. (E) HNF4A ChIP-Seq showing its binding peak at the enhancer of IL4R in Eso26 (EAC), Caco2 (COAD), GP5d (COAD) and LoVo (COAD) cells. (F) qRT-PCR analysis showed that knockdown of HNF4A decreased the expression level of the HNF4A targets of Interleukin signaling. (G) IL4R, LYN, ELK1 and IL6ST was silenced by individual siRNAs in Eso26 cells, and cell proliferation was measured. Mean ± s.d. are shown. *, P<0.05; **, P<0.01.

### HNF4A transcriptionally activates Interleukin signaling pathway in GIAC cells

To investigate the downstream targets of HNF4A in GIAC cells, we performed RNA-seq upon knockdown of this MRTF in ESO26 cells. Differential expression analysis of replicates identified 335 up-regulated genes and 346 down-regulated genes when compared with scrambled control (fold change > 2, p < 0.05). Pathway enrichment analysis of the differential expressed genes showed that a number of cancer-related pathways were top ranked, such as Interleukin signaling, WNT signaling and MAPK1/3 pathway (Fig. 7D). Among these, Interleukin signaling pathways were most notable, as they were the top 3 most significantly enriched. Specifically, a total of 21 genes of this pathway were downregulated upon knockdown of HNF4A. Integrating HNF4A ChIP-Seq results found that 14 of the 21 pathway components were directly bound by HNF4A (Fig. 7E and **Supplementary Table 2**). We randomly selected a number of targets and RT-PCR analysis confirmed their expression changes (Fig. 7F). Some of these 14 direct targets (particularly ELK1, LYN, IL4R and IL6ST) have a pro-tumor properties in several cancer types[35–41] but their functions have not been explored in GIAC. We thus next tested whether they contributed to HNF4A-mediated oncogenic functions in GIAC cells. Importantly, silencing any of these 4 genes significantly inhibited the proliferation of GIAC cells (Fig. 7G). Together with our earlier result that HNF4A regulated HNF1A (another candidate GIAC MRTF, which also promoted GIAC cell proliferations, **Supplementary Figs. 6A,B**), these data demonstrate that HNF4A plays an important role in promoting GIAC proliferation and survival by transcriptionally activating many downstream targets, including HNF1A and those belonging to the Interleukin signaling (Fig. 8).

**Figure 8.**
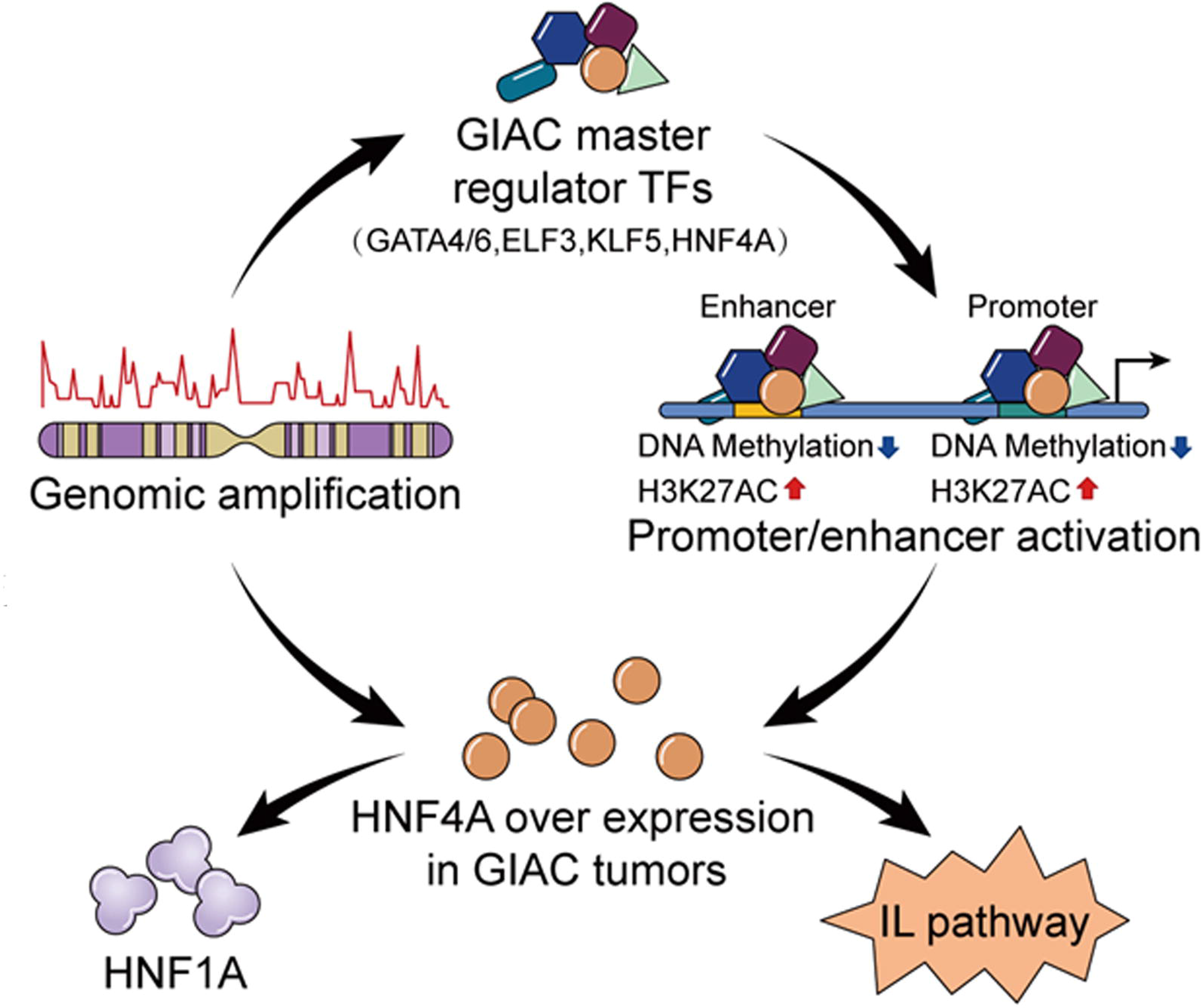
Proposed model of epigenomic and genomic activation of lineage-specific oncogene HNF4A which promotes GIAC cancers.

## Discussion

The clinical management of GIACs has advanced only modestly over the last several decades, and thus urgent needs exist to decipher the biological and pathological basis of these malignancies to improve prevention, diagnosis and therapy. Recently, unbiased genomic studies have strongly suggested unified genomic features among different types of GIACs, which has prognostic and therapeutic implications. For example, MSI-high STAD and COAD patients have shown better clinical response to immune checkpoint blockade independent of anatomic tumor origin[5, 6]. In addition, ERBB2 amplified and overexpressed EAC, STAD and COAD patients similarly benefit from anti-Her2 therapy. Thus, identification and characterization of unified genomic and/or epigenomic features across different types of GIACs may have important clinical implications. In this study, we aimed to identify such shared epigenomic characteristics across anatomically distinct GIACs. Motivated by the prominent changes in DNA methylation and gene expression in cell type- and cancer-specificity manner, we inferred gene expression networks, cis-regulatory elements as well as their upstream MRTFs in different types of GIACs using a novel mathematical tool (ELMER) we developed previously. This algorithm provides an unbiased systematic approach to reconstruct gene regulatory networks by integrating matched methylome and gene expression data.

We found that different types of GIACs display notable similarity with respect to their hyperactive upstream MRTFs, which in turn conceivably orchestrate similar downstream transcriptional programs. Indeed, some of the MRTFs (such as GATA4/6, KLF5 and CDX2) are known to have similar oncogenic functions in different types of GIACs. As ELMER is based on DNA methylation array data which has limited resolution and representability, we generated WGBS results in an independent cohort of EAC and ESCC samples and successfully validated the ELMER prediction. These results highlight that our approach is capable of identifying *bona fide* MRTFs for cancer biology, and the list of candidate GIAC MRTFs warrant further investigation.

Focusing on the most significant MRTF, HNF4A, we first observed that the genomic and epigenomic activation of this factor was unique and specific in GIACs. Specifically, Pan-Cancer transcriptomic data from either TCGA or CCLE showed that the expression of HNF4A was uniquely high in GIACs comparing with other cancer types. Both internal IHC staining of different GIAC samples and Human Protein Atlas results consistently identified the same lineage-specific pattern of HNF4A protein. Moreover, higher HNF4A expression was associated with worse overall survival in different types of GIAC patients, supporting a shared functional contribution to GIAC biology independent of anatomic origin. Genomic analyses demonstrated that copy number amplification accounted partially for the GIAC-specific over-expression of HNF4A. The other important source to drive the high HNF4A level was epigenomic activation, which was supported by ATAC-Seq, WGBS as well as H3K27ac ChIP-Seq profiling. These epigenomic analyses identified that GIAC-specific open chromatin regions, which were concordantly de-methylated and had high H3K27ac signals, likely contributed to the transcriptional activation of HNF4A in GIAC in a lineage-specific manner. Prompted by this epigenomic observation, we next characterized the underlying mechanisms. We first applied 4C-Seq and identified three enhancer elements (E1-E3) interacting with the *HNF4A* promoter. Motif enrichment analysis of these regulatory regions coupled with ChIP-Seq experiments identified that they were occupied by five upstream GIAC-specific MRTFs, GATA4/6, ELF3, KLF5 and HNF4A itself. Luciferase reporter assays and loss-of-function experiments of individual MRTFs confirmed that HNF4A transcription is activated by these five MRTFs through interacting with HNF4A promoter and three local enhancers. HNF4A has been shown to be regulated by GATA4, GATA6 and KLF5 in STAD cells[10], which is consistent with our unbiased motif search and epigenomic profiling. Furthermore, the self-regulatory property of HNF4A identified in the present study has been observed in many TFs in different cell types, which is regarded as an important and common characteristic of key MRTFs[32, 42].

Functionally, HNF4A plays a role in normal development of the liver[43–45], kidney[46], and intestine[47, 48]. In cancer biology, HNF4A has opposite roles in different cell types. For example, HNF4A is a tumor suppressor in hepatocellular cancer[49] while it was required for maintaining the proliferation of STAD cells[10, 28, 29]. During the preparation of the manuscript, another study was published suggesting that HNF4A-mediated transcriptional program was more active in both Barrett’s esophagus and EAC cells than normal esophageal squamous cells[50]. Using analysis from unbiased shRNA library screen of over 500 cancer cell lines, we showed that HNF4A is uniquely and specifically essential for the viability of GIAC cells. We further confirmed this finding using both genetic and chemical tools in different types of GIAC cells in vitro and in vivo, highlighting HNF4A as a prominent lineage-specific oncogene in GIAC tumors. Interestingly, a previous report showed that HNF4A activity could be inhibited by metformin[28], making HNF4A an appealing and specific therapeutic target for GIAC, including early chemoprevention. We further performed ChIP-Seq and transcriptomic analysis and identified Interleukin signaling as a key downstream pathway of HNF4A. Considering that agents inhibiting Interleukin signaling (such as Secukinumab, Ustekinumab and Ixekizumab) are available clinically, these results together highlight HNF4A-Interleukin axis as a potential actionable cascade for GIAC patients.

## Conclusion

In summary, applying ELMER to matched DNA methylation and expression profiles to reconstruct TF networks, the present work identifies a panel of hyperactive MRTFs shared by different GIAC cancers. As a top candidate, HNF4A is highlighted as a key oncogenic MRTF with prominent genomic and epigenomic activations in GIAC-specific manner. By providing mechanistic insights into the upstream and downstream regulation of HNF4A, this work significantly advances our understanding of the GIAC-specific gene regulatory networks, while providing potential therapeutic strategies against these common cancers.

## Methods

We applied mathematical algorithms to large-scale DNA methylome and transcriptome profiles to reconstruct transcription factor (TF) networks using 907 GIAC samples from The Cancer Genome Atlas (TCGA). Complementary epigenomic technologies were performed to investigate HNF4A activation, including Circularized Chromosome Conformation Capture (4C), Chromatin immunoprecipitation (ChIP) sequencing, Whole Genome Bisulfite Sequencing (WGBS), and Assay for Transposase-Accessible Chromatin (ATAC) sequencing. In vitro and in vivo cellular phenotypical assays were conducted to study HNF4A functions.

### Antibodies and reagents

The following antibodies and reagents were used: Anti-HNF4A for ChIP-Seq and WB (Abcam, ab41898), Anti-HNF4A for IHC (a gift from Kenji Daigo and Takao Hamakubo), Anti-ELF3 (Santa Cruz Biotechnology, sc-376055 X), Anti-KLF5 (Santa Cruz Biotechnology, sc-398470 X), Anti-GATA6 (Cell Signaling Technology, 5851), Anti-GATA4 (Thermo Fisher Scientific, MA5-15532), Anti-HNF1A (Cell Signaling Technology,89670), Anti-FLAG (Sigma, F1804), Anti-Actin (Santa Cruz Biotechnology, sc-8432), Anti-H3K27Ac (Abcam, ab4729), Anti-mouse IgG-HRP (Jackson ImmunoResearch Laboratories, Inc., 115-035-003), Anti-rabbit IgG-HRP (Jackson ImmunoResearch Laboratories, Inc., 115-035-144), FITC Annexin V Apoptosis Detection Kit (BD Biosciences, 556547), Lipofectamine RNAiMAX (Thermo Fisher, 13778150), BI-6015 (Cayman, 12032) and siRNAs were purchased from Suzhou GenePharma Co., Ltd. (Sequences are provided in Supplementary Table 1).

### Human cancer cell lines

Eso26, OACM5.1, SNU398 cells were grown in RPMI-1640 medium, and SKGT4, ASPC-1, Suit2, colo205 were cultured in Dulbecco’s modified Eagle medium (DMEM). Eso26 and OACM5.1 were purchased from the American Type Culture Collection (ATCC). ASPC-1, Suit2, colo205 and SNU398 were purchased from JENNIO Biological Technology (Guangzhou, China). Both media were supplemented with 10% fetal bovine serum (Omega Scientific, Tarzna, CA), 100 U/ml penicillin and 100 mg/ml streptomycin. All cell lines were verified by short tandem repeat analysis in the year of 2018.

### ELMER analyses

Illumina HM450 methylation data from 4 different TCGA projects, COAD-READ, ESCA, PAAD, and STAD were processed with SeSAMe[51], and the matched RNA-seq (FPKM-UQ) data were downloaded from GDC (Genomic Data Commons, https://portal.gdc.cancer.gov/) from the harmonized database (data aligned to hg38/GRCh38). Unsupervised analysis was performed using ELMER[52] for the following pair-wise comparisons: EAC primary tumors (n = 78) vs ESCC primary tumors (n = 76); EAC primary tumors (n = 78) vs EAC normal adjacent samples (n = 5), COAD-READ primary tumors (n = 389) vs colon normal adjacent samples (n = 21), PAAD primary tumors (n = 177) vs pancreas normal adjacent samples (n = 4). Because of the small number of normal adjacent samples from STAD (n=2) which also lacked RNA-seq data, we could not perform comparison for STAD primary tumors. We applied the default probe filter “distal”, which uses 160,944 probes that are > 2 kbp from any transcription start site as annotated by GENCODE 28. ELMER version 2.5.4 was used with the following parameters: get.diff.meth(sig.diff) = 0.3, get.diff.meth(p_value) = 0.01, get.diff.meth (minSubgroupFrac) = 0.2, get.pair(Pe) = 10^-3, get.pair(raw.pvalue) = 10^-3, and get.pair(filter.probes) = TRUE, get.pair(permu.size) = 10000, get.pair(minSubgroupFrac) = 0.4, get.enriched.motif(lower.OR) = 1.1, get.TFs(minSubgroupFrac) = 0.4. For each comparison, the TF subfamilies were inferred from the TFs which had the most significant anti-correlation scores. The candidate MRTFs were next identified within the TF subfamily with FDR < 0.05 using the classification from TFClass[53].

### Whole genome bisulfite sequencing (WGBS) and data analysis

Primary tumor tissues from 5 EAC and 12 ESCC samples were collected at Shantou Center Hospital, China. All patients were consented for molecular analysis. This study has been approved by the ethics committee of the Medical College of Shantou University. WGBS library preparation and sequencing were performed by Novogene, Inc. Briefly, 2-3 ug DNA spiked with 26 ng lambda DNA were fragmented. Cytosine-methylated barcodes were ligated to the sonicated DNA, treated twice with bisulfite using EZ DNA Methylation-GoldTM Kit (Zymo Research). The resulting DNA fragments were PCR amplified using HiFi HotStart Uracil + ReadyMix (Kapa Biosystems). The clustering of the index-coded samples was performed on Illumina cBot Cluster Generation System according to the manufacturer’s instructions. DNA libraries were sequenced on Illumina Hiseq platform with 150bp paired-end reads.

WGBS reads were aligned to the genome (build GRCh38) using BISCUIT (https://github.com/zwdzwd/biscuit). Duplicated reads were marked using Picard Tools (https://broadinstitute.github.io/picard/). Methylation rates were called using BISCUIT. CpGs with fewer than 5 reads of coverage were excluded from further analysis. Quality control was performed using TrimGalore by default parameter for Illumina sequencing platforms, (https://www.bioinformatics.babraham.ac.uk/projects/trim_galore/), PicardTools as well as MultiQC (https://multiqc.info/). Bisulfite non-conversion was checked using the Biscuit QC module in MultiQC (https://github.com/ewels/MultiQC/tree/master/multiqc/modules/biscuit).

### ELMER-dependent motif analysis of WGBS

EAC and ESCC WGBS BED files were converted into Tag Directories using HOMER (http://homer.ucsd.edu/homer/) with the mCpGbed option. The annotatePeaks.pl script from HOMER was then run on all Tag Directories from each EAC and ESCC sample, with focal points defined as the list of predicted HNF4A motifs adjacent to EAC-demethylated CpGs, generated by ELMER (EAC vs. ESCC run described above). A bin size of 100bp for HOMER to generate spatial methylation plot.

### ELMER-independent motif analysis of WGBS

The WGBS analysis was performed as described in the ELMER-dependent section above, except with different focal points. Here, we generated a set of predicted HNF4A binding sites by scanning the complete human genome using HOMER’s scanMotifGenomeWide.pl at a p-value threshold of 1E-5. We used the same HOCOMOCO[54] HNF4A model as used in ELMER (http://hocomoco11.autosome.ru/motif/HNF4A_HUMAN.H11MO.0.A). We then filtered only those predicted HNF4A sites that overlapped one of the 72,264 esophageal-specific ATAC-seq peaks from TCGA[25], aligning to the HNF4A motif for the spatial methylation plot. For the box plot, methylation was averaged within 20kb from each HNF4A motif.

### CCLE data

We used dependency scores from the “Combined RNAi (Broad, Novartis, Marcotte)” field, expression values from the “Expression public 19Q2” field, and methylation values from the “Methylation (1kb upstream TSS)” field.

### Chromatin immunoprecipitation (ChIP) sequencing

1×10^7^∼ 5×10^7^cells were harvested in 15 ml tubes and fixed with 2 ml of 1% paraformaldehyde for 10 min at room temperature, which was stopped by 2 ml of 250 mM of glycine. Samples were rinsed with 1xPBS twice and lysed twice with 1 ml of 1 X lysis/wash buffer (150 mM NaCl, 0.5 M EDTA pH 7.5, 1M Tris pH 7.5, 0.5% NP-40). Samples were filtered through a 29 G needle during each lysis process, and were harvested by centrifuge. Cell pellets were resuspended in sharing buffer (1% SDS, 10 mM EDTA pH 8.0, 50 nM Tris pH 8.0) and sonicated in a Covaris sonicator. The sonicated samples were subsequently centrifuged to remove debris and supernatants were diluted 5X with dilution buffer (0.01% SDS, 1% Triton X-100, 1.2 mM EDTA pH 8.0, 150 nM NaCl). The primary antibodies were then added and incubated by rotation at 4_J overnight. Dynabeads Protein G beads (Life Technologies) were added the next morning and incubated by rotation for 4 hours. The beads were washed with 1x lysis/wash buffer followed by wash in cold TE buffer. DNA molecules were reverse crosslinked, purified and subject to library preparation and sequencing on Illumina HiSeq platform.

### Immunohistochemistry (IHC) staining and quantification

Tissue microarrays were purchased from US Biomax Company (Col05-118e, ES8011a, PA804a). All slides were deparaffinized and immersed before incubating with anti-HNF4A antibody. IHC staining was performed at room temperature using a PowerVision Homo-Mouse IHC kit (ImmunoVision Technologies, Daly City, CA). Hematoxylin (Sigma) was used for counterstaining. HNF4A immunopositivity was scored as follows: 0, no staining or sporadic staining in < 5% cells; 1, weak staining in 5–25% cells; 2, weak staining in 26–50% of tumor cells; 3, strong staining in 26–50% cells; and 4, strong staining in > 50% cells.

### Xenograft assays in nude mice

For shRNA-based experiments, 6 male six-week-old mice were randomly separated to two groups and subcutaneously injected with 2×106 Eso26 cells expressing inducible scrambled shRNA or shRNA against HNF4A. After tumor inoculation, all mice were fed with 2mg/ml doxycycline (Abcam, ab141091) containing water. For the BI-6015 experiment, 6 male six-week-old mice were subcutaneously injected with 2×106 Eso26 cells. Either BI-6015 or vehicle control (5% DMSO+45% PEG 300+H2O) were administered three times per week by intraperitoneal injection at either 50 mg/kg or 100 mg/kg. Xenograft size was measured two times per week for a total of 4 weeks. Mice were euthanized at the end of experiment and xenograft tumors were extracted for analysis.

### Construction of expression vectors

The pBABE-puro-HNF4A expression vector was amplified based on CMV-68 HNF4A (Addgene, #31092) and a 3xFLAG-tag was added via PCR. The amplified 3xFLAG-tagged HNF4A was then cloned into pBABE-puro vector (Addgene, #1764). The shRNA expression vector was designed based on siRNA sequences (provided in Supplementary Table 1) and cloned into Tet-pLKO-puro vector (Addgene, #21915) and pLKO-puro vector (Addgene, #8453). To produce viral particles, recombinant viral vectors and packaging vectors were co-transfected into 293T cells. Supernatants containing viral particles were harvested and filtered through a 0.45 µM filter 48 hours after transfection. GIAC cells were then infected with the virus in the presence of 10 mg/ml Polybrene.

### In vitro cell proliferation assay

3,000 - 5,000 GIAC cells were seeded into 96-well plates and cultured for indicated periods of time. For inhibitor treatment, culture medium was replaced with either DMSO- or BI-6015-containing medium after 24 hours. Cell proliferation was measured by staining of 3-(4,5-dimethylthiazol-2-yl)-2,5-diphenyltetrazolium bromide (MTT).

### Luciferase reporter assay

Three enhancer elements (E1: chr20:42,838,300-42,839,699, E2: chr20:42,931,956-42,930,286, E3: chr20: 43,035,000-43,037,051 and negative control chr20:42,860,825-42,862,318) were cloned into pGL3-Promoter luciferase reporter vector (Promega). HNF4A promoter luciferase reporter vector HNF4A-P2-2200 was purchased from Addgene (Addgene, 31062). HNF1A promoter luciferase reporter vector HNF1A-P-1526 was cloned in house. A Renilla luciferase control vector was co-transfected for normalization. After 48 hours of transfection, the reporter activity was measured by the Dual-Luciferase Reporter Assay System (Promega, E1910).

### Preparation of 4C-Seq library

4C-Seq libraries were prepared using our previous described protocols[55]. In brief, Eso26 cells were single-cell suspended, and the chromatin was cross-linked with 1% formaldehyde at room temperature. Cells were lysed and DNA was digested with the restriction enzyme TaqI (R0149L, NEB). Digested DNA was next ligated using T4 DNA ligase (EL0013, Thermo Scientific), followed by removal of cross-link using Proteinase K (19133, Qiagen), yielding 3C libraries. These 3C libraries were subjected to a second enzyme digestion by Csp6i (R0639L, NEB), followed by another ligation using T4 DNA ligase. A total of 3.2 µg of the resulting 4C templates was used for a scale-up inverse nested PCR for 29 cycles. The PCR products were purified using Macherey Nagel Nucleospin Gel and PCR Purification kit (Takara Bio). 4C sequencing libraries were made from the PCR products using Thruplex DNA-seq kit (R400427, Takara Bio). The libraries were subjected to Agencourt AMPure XP Bead clean-up (A63881, Beckman Coulter) using a bead-to-DNA ratio of 1:1. Libraries were sequenced using 1 × 75bp for 2 million read depth using MiniSeq system (Illumina).

### Primer design for 4C-Seq

The inverse primers were designed based on a viewpoint region. DpnII restriction sites flanking the region of interest were identified and the sequence between the nearest DpnII and TaqI restriction sites were selected as the viewpoint region. Based on this region, 4C primers (F1-AGGAGACGGACCTTAATCAGATC, R1-AATGTGACCGCTTCCCTAAGCT) were designed with the following settings: optimal melting temperature of 60 _JC (57 _JC - 62 _JC); GC content: 40 - 60%.

### 4C-Seq data analysis

4C sequencing data were analyzed using the R package r3CSeq[56]. Briefly, for each replicate the raw reads were aligned to the reference human genome which was masked for the gap, repetitive and ambiguous sequences. The masked version of the genome was downloaded from the R Bioconductor repository (BSgenome.Hsapiens.UCSC.hg19.masked). The number of mapped reads for each window were counted and normalized to obtain RPM (reads per million per window) values, which was used to perform statistical analysis. Interacted regions were plotted and visualized on UCSC Genome Browser as custom tracks. 4C sequencing data have been deposited to Gene Expression Omnibus (GSE132813, token: ehixwcucnjoxvsb).

### ChIP-Seq data analysis

We generated ChIP-Seq data for HNF4A and GATA4 in Eso26 cells in the present work, which has been deposited to Gene Expression Omnibus (GSE132813, token: ehixwcucnjoxvsb). ChIP-Seq data of GATA6, KLF5 and ELF3 in Eso26 cells, and H3K27Ac ChIP-Seq data in the following cell lines were generated in-house recently (GSE132686 and GSE106563) [57, 58]: ESO26, Flo1, JH, KYAE1, OACM5.1, OACp4C, OE19, OE33, SKGT4, KYSE70, KYSE140, KYSE510, TT, TE5 and TE7. H3K27Ac ChIP-Seq from STAD normal (n=20) and tumor samples (n=20) were from Wen Fong Ooi et al., 2016[59]. For data analysis, ChIP-Seq reads were aligned to reference genome (UCSC hg19) using Bowtie (v1.1.2)[60] or BWA[61] under default parameters. PCR duplicates were removed using Picard tools. Peaks were filtered to remove all peaks overlapping ENCODE blacklisted regions. The bigwig files were generated by bamCompare in DeepTools (v3.1.3)[62].

Motif enrichment analyses were performed using bedtools (v2.27.1)[63] and HOMER (v4.10.3)[64]. Briefly, genomic regions of E1-E3 and HNF4A promoter were divided into 557 sub-region (25bp each) using makewindows function in bedtools. The resultant file containing 557 sub-region were used to identify enriched motifs using HOMER with the parameters -size 200 -lens 8,10,12.

## Abbreviations

Esophageal squamous cell carcinoma (ESCC);Esophageal adenocarcinoma (EAC);Transcription factor (TF);Chromatin immunoprecipitation sequencing (ChIP-Seq);Histone 3 lysine 27 acetylation (H3K27Ac);The Cancer Genome Atlas (TCGA);Cancer Cell Line Encyclopedia (CCLE);Circularized chromosome conformation capture (4C);Fluorescence-activated cell sorting (FACS);Immunohistochemistry (IHC).

## Declarations

### Availability of data and materials

4C sequencing data have been deposited to Gene Expression Omnibus (GSE132813, token: ehixwcucnjoxvsb). ChIP-Seq data for HNF4A and GATA4 in Eso26 cells in the present work, which has been deposited to Gene Expression Omnibus (GSE132813, token: ehixwcucnjoxvsb). ChIP-Seq data of GATA6, KLF5 and ELF3 in Eso26 cells, and H3K27Ac ChIP-Seq data in the following cell lines were generated in-house recently (GSE132686 and GSE106563). To whom correspondence and material requests: De-Chen Lin, PhD (Phone: +1-310-423-7740).

## Author Contributions

D.-C.L. conceived and devised the study. J.P., T.C.S., N.G., Q.Y., J.P., B.P.B. and D.-C.L. designed experiments and analysis. J.P., T.C.S., N.G., Q.Y., J.P., S.C., L.-W.D. and Y.-Y.J. performed the experiments. T.C.S., N.G., Q.Y. and B.P.B. performed bioinformatics and statistical analysis. S.-Y.H., L.-Y.X., E.-M.L., S.G, K.D., T.H., Y.B.D. and S.J.K. contributed reagents and materials. J.P., T.C.S., B.P.B., H.P.K. and D.-C.L. analyzed the data. B.P.B., H.P.K. and D.-C.L. supervised the research and all authors wrote the manuscript.

