## Supplemental Figures for "Lineage-Specific Epigenomic and Genomic Activation of the Oncogene HNF4A Promotes Gastrointestinal Adenocarcinomas"

**Supplementary Figures & Legends**

**Supplementary Figure 1**

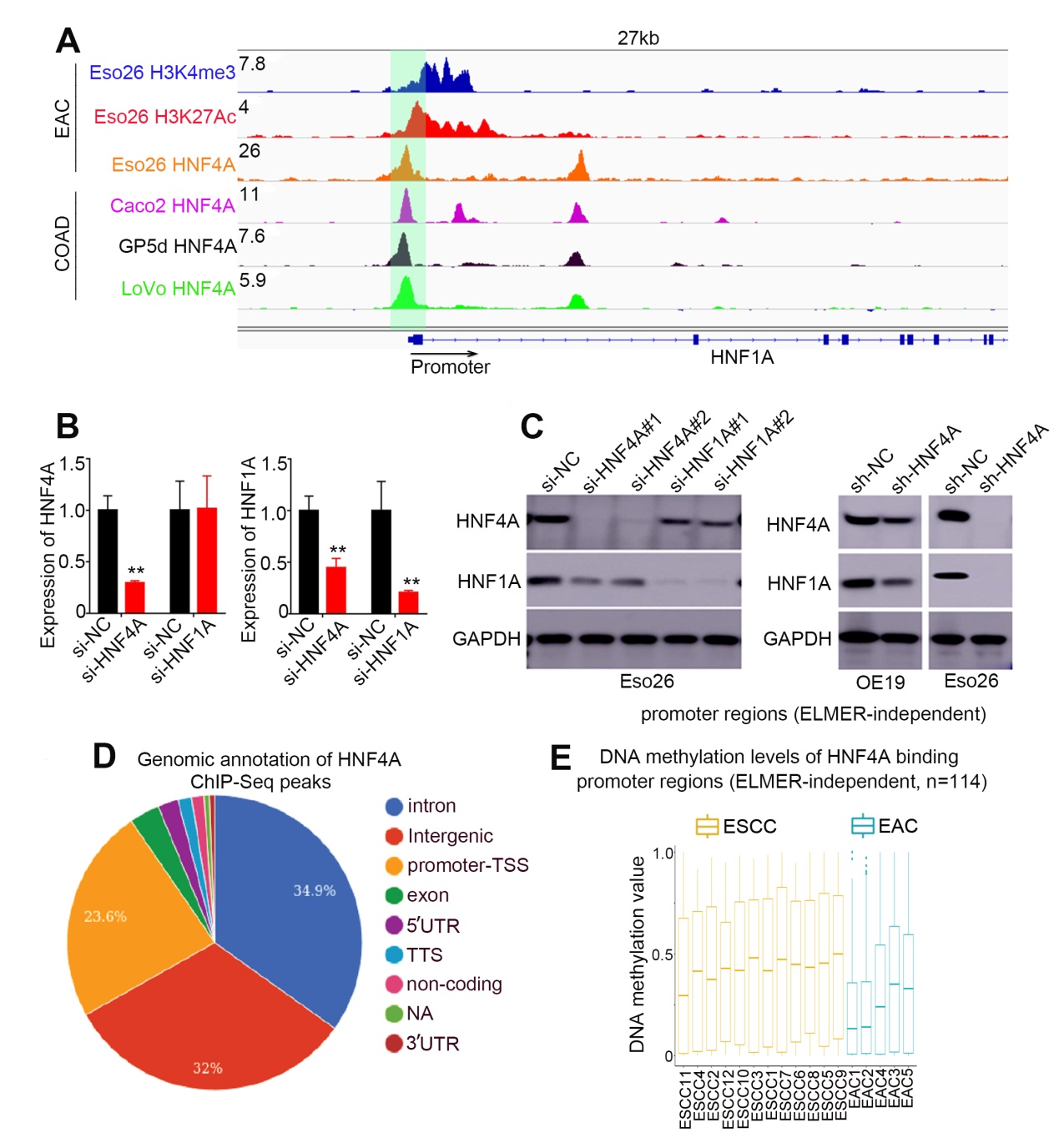

**Figure S1. HNF4A regulates the promoter of HNF1A.** (A) HNF4A ChIP-Seq showing its binding peak at the promoter region of HNF1A in Eso26 (EAC), Caco2 (COAD), GP5d (COAD) and LoVo (COAD) cells. (B) qRT-PCR and (C) Western Blotting analyses showed that knockdown of HNF4A decreased the expression level of the HNF1A while the opposite was not observed. (D) Genomic distribution of HNF4A binding sites from HNF4A Chip-seq data in ESO26 cells. (E) Box plots showing the methylation level of ELMER-independent HNF4A-binding promoter regions using in-house WGBS data.

**Supplementary Figure 2**

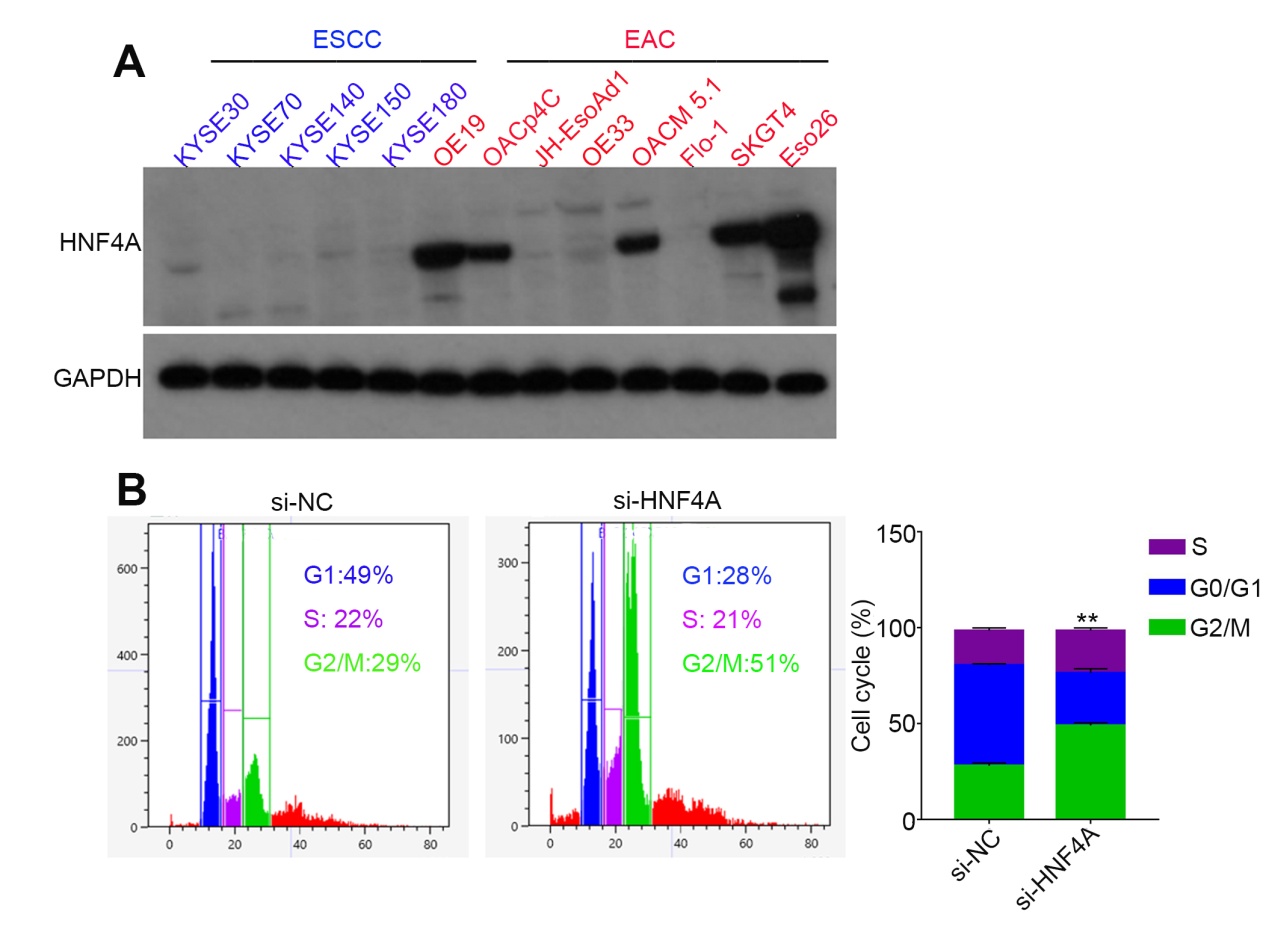

**Figure S2. HNF4A regulates the cell cycle and apoptosis of GIAC cells.** (A) Western Blotting analyses HNF4A in ESCC and EAC cell lines. (B) Cell cycle assays of Eso26 cells upon knockdown of HNF4A with siRNA.

**Supplementary Figure 3**

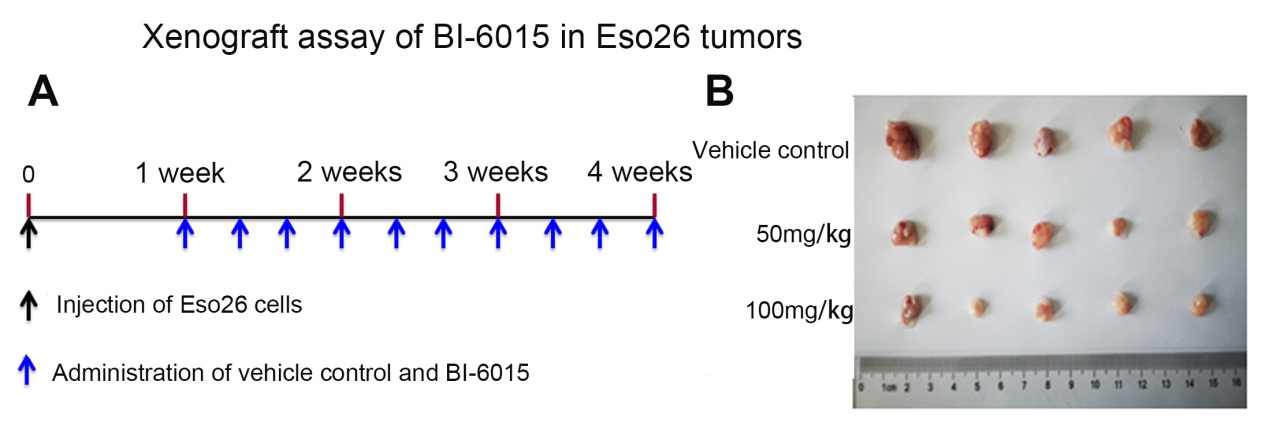

**Figure S3. Targeting HNF4A with BI-6015 in GIAC cells *in vivo*.** (A)Injection time and dose of BI-6015 and vehicle control (5% DMSO+45% PEG 300+H2O). (B) Mice were euthanized at the end of experiment and xenografts tumor were significantly inhibited by BI-6015.

**Supplementary Figure 4**

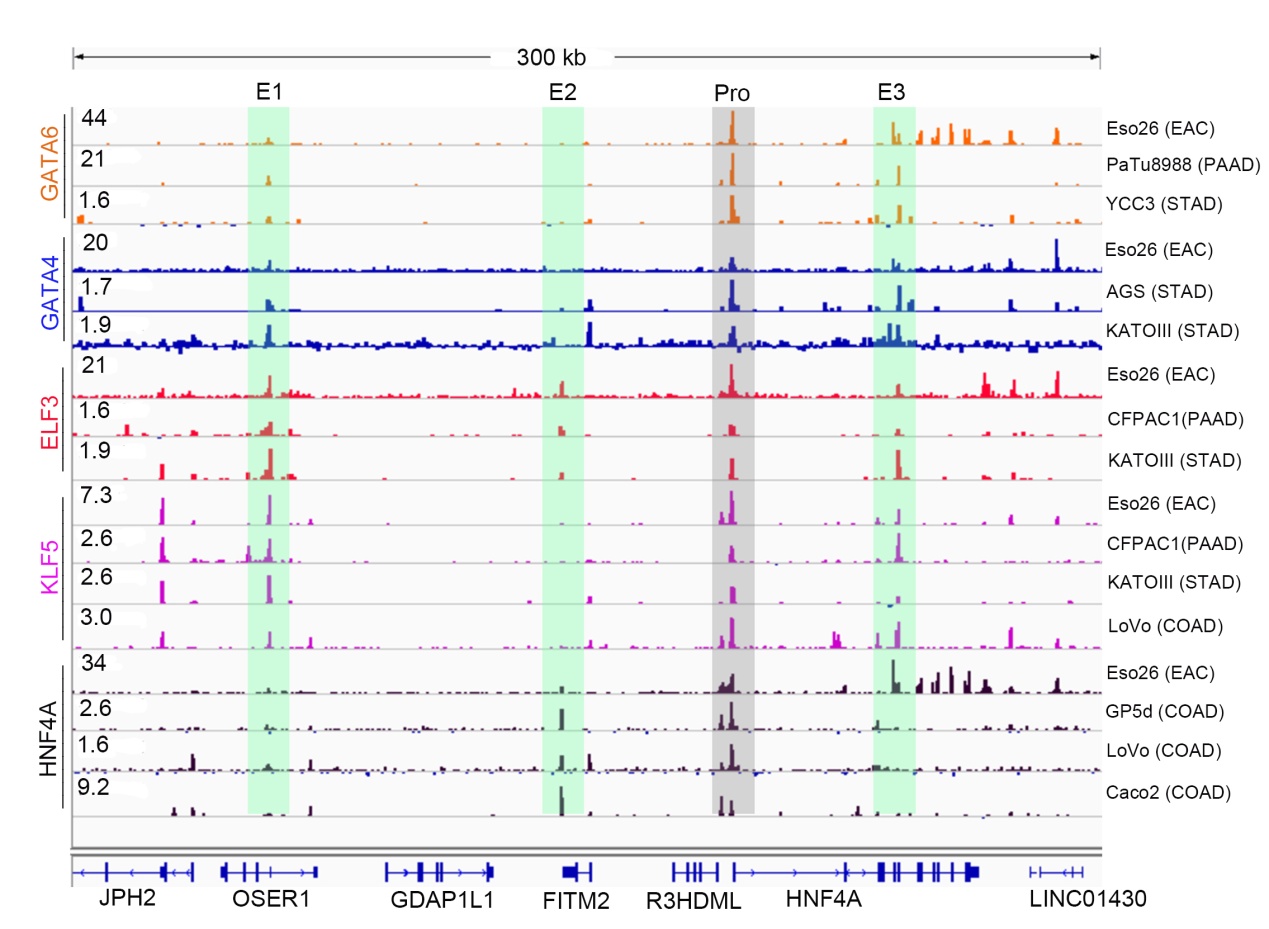

**Figure S4. MRTFs occupy and regulate HNF4A promoter and enhancers in GIAC cells.** ChIP-Seq profiles for indicated MRTFs at HNF4A promoter and enhancers loci in GIAC cells.

**Supplementary Figure 5**

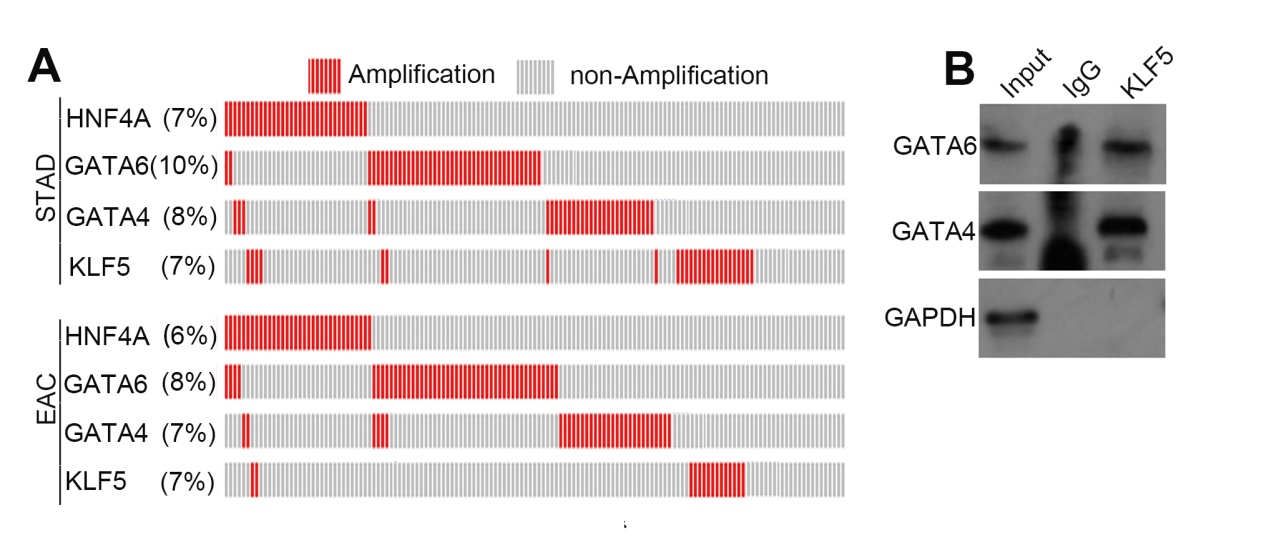

**Figure S5. Upstream regulation of HNF4A transcription by GIAC master TFs.** (A) Copy number gains of HNF4A, GATA4/6 and KLF5 in TCGA EAC and STAD cohorts. Each bar represents one sample. (B) Co-IP assay of GATA4, GATA6 and KLF5 in Eso26 cells.

**Supplementary Figure 6**

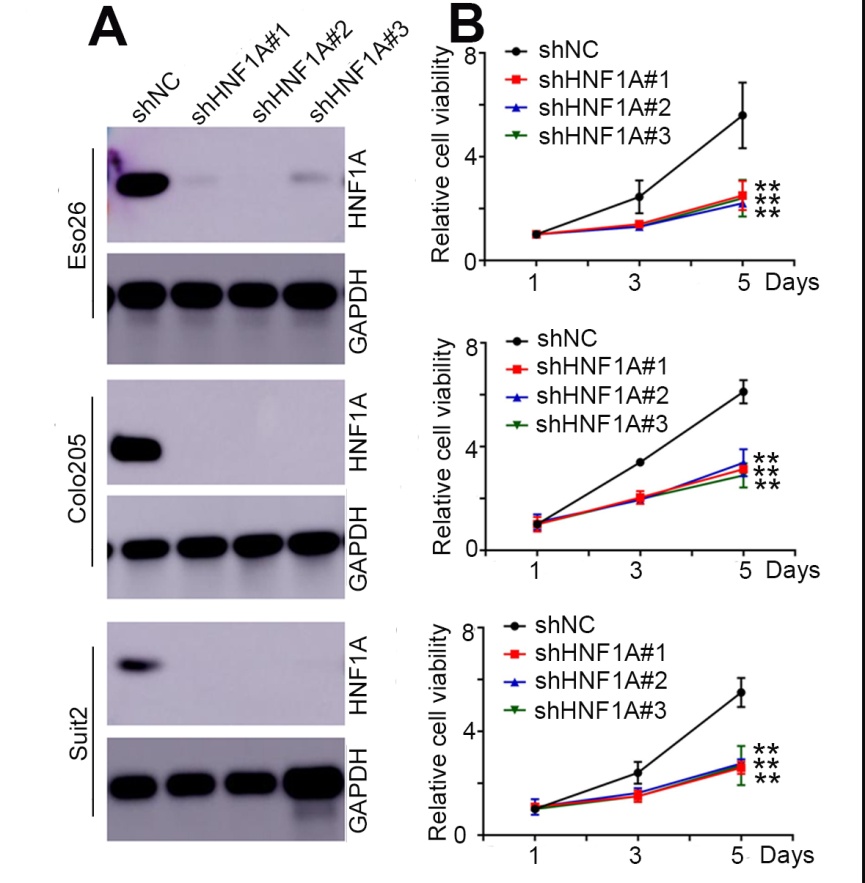

**Figure S6. HNF1A promotes the proliferation and survival of GIAC cells.** (A) In EAC (Eso26), COAD (Colo205) and PAAD (Suit2) cell lines, HNF4A expression was silenced by three different shRNAs and followed by Western Blotting assay. (B) Cell proliferation assays in GIAC cells. Mean ± s.d. are shown. *, P<0.05; **, P<0.01.
